# Mating system is associated with seed phenotypes upon loss of RNA-directed DNA Methylation in Brassicaceae

**DOI:** 10.1101/2023.07.03.547469

**Authors:** Kelly J. Dew-Budd, Hiu Tung Chow, Timmy Kendall, Brandon C. David, James A. Rozelle, Rebecca A. Mosher, Mark A. Beilstein

## Abstract

In plants, *de novo* DNA methylation is guided by 24-nt short interfering (si)RNAs in a process called RNA-directed DNA methylation (RdDM). Primarily targeted at transposons, RdDM causes transcriptional silencing and can indirectly influence expression of neighboring genes. During reproduction, a small number of siRNA loci are dramatically upregulated in the maternally-derived seed coat, suggesting that RdDM might have a special function during reproduction. However, the developmental consequence of RdDM has been difficult to dissect because disruption of RdDM does not result in overt phenotypes in *Arabidopsis thaliana*, where the pathway has been most thoroughly studied. In contrast, *Brassica rapa* mutants lacking RdDM have a severe seed production defect, which is determined by the maternal sporophytic genotype. To explore the factors that underlie the different phenotypes of these species, we produced RdDM mutations in three additional members of the Brassicaceae family: *Camelina sativa, Capsella rubella*, and *Capsella grandiflora*. Among these three species, only mutations in the obligate outcrosser, *C. grandiflora*, displayed a seed production defect similar to *Brassica rapa* mutants, suggesting that mating system is a key determinant for reproductive phenotypes in RdDM mutants.

## Introduction

RNA-directed DNA methylation (RdDM) triggers de novo DNA methylation and maintains non-symmetric methylation at euchromatic transposons (Matzke and Mosher, 2014; Erdmann and Picard, 2020). This methylation causes transcriptional silencing to prevent the mobilization of transposons and can indirectly influence expression of neighboring genes (Hollister and Gaut, 2009; Hollister et al., 2011; Forestan et al., 2017). RdDM employs 24-nt small interfering (si)RNAs, which are produced though the sequential activities of RNA Polymerase IV (Pol IV), RNA-DEPENDENT RNA POLYMERASE2 (RDR2), and DICER-LIKE3 (Singh et al., 2019; Fukudome et al., 2021; Huang et al., 2021; Wang et al., 2021; Loffer et al., 2022). These siRNAs are loaded into ARGONAUTE4 and direct the protein to nascent transcripts produced by RNA Polymerase V (Wierzbicki et al., 2009; Liu et al., 2018; Singh et al., 2019; Wang et al., 2023). Once assembled at the chromatin, ARGONAUTE4 recruits DOMAINS REARRANGED METHYLTRANSFERASE to methylate DNA proximal to the Pol V transcript (Zemach et al., 2013; Stroud et al., 2014; Zhong et al., 2014). Because non-symmetric DNA methylation is not maintained on both daughter strands during DNA replication, repeated *de novo* methylation is necessary to maintain DNA methylation (Law and Jacobsen, 2010; Law et al., 2013).

Twenty-four nt siRNAs are particularly abundant in reproductive tissues, suggesting that RdDM has a special role during reproduction (Chow and Mosher, 2023). Increased accumulation of 24-nt siRNAs is due to expression of reproductive-specific loci, rather than upregulation of all RdDM sites (Mosher et al., 2009; Rodrigues et al., 2013; Grover et al., 2020; Li et al., 2020; Long et al., 2021; Zhou et al., 2022). In particular, ovule- and endosperm-specific siRNA loci have been described in rice (Rodrigues et al., 2013), *Brassica rapa* (Grover et al., 2020), and Arabidopsis (Zhou et al., 2022). Known as siren loci, these siRNAs comprise approximately 90% of the siRNA population in unfertilized ovules and trigger methylation of protein-coding genes during seed development (Grover et al., 2020; Burgess et al., 2022). In some cases, siren-induced *trans-*methylation impacts gene expression (Burgess et al., 2022).

The developmental consequence of RdDM action during reproduction has been difficult to dissect because in *Arabidopsis thaliana*, where the pathway has been most thoroughly studied, disruption of RdDM does not result in overt phenotypes. There is a slight reduction in seed weight for homozygous mutant seed (Grover et al., 2018) and an increase in the success of paternal-excess crosses when RdDM is eliminated from the paternal parent (Erdmann et al., 2017; Martinez et al., 2018; Satyaki and Gehring, 2019). In contrast, *Brassica rapa* RdDM mutants are almost completely sterile and produce seeds that are smaller than wild type (Grover et al., 2018). Reproductive defects are also among the pleiotropic phenotypes observed in RdDM mutants of maize, tomato, and rice (Erhard et al., 2009; Gouil and Baulcombe, 2016; Xu et al., 2020; Zheng et al., 2021; Chakraborty et al., 2022; Wang et al., 2022).

The different phenotypes in *A. thaliana* and *B. rapa* raises the question: what factors underlie the phenotypic differences upon loss of RdDM in these two relatively closely related species of Brassicaceae? It has been proposed that RdDM might play a larger developmental role in species with a higher proportion of transposons, or in genomes where transposons are in closer proximity to protein-coding genes, as these factors increase the possibility that transcriptional silencing of a transposon will have an indirect impact on a protein-coding gene (Wei et al., 2014; Gouil and Baulcombe, 2016; Wang et al., 2020). We propose two additional hypotheses to explain the different phenotypes of *A. thaliana* and *B. rapa* RdDM mutants.

Firstly, RdDM might play a role in mediating conflicts among the subgenomes of polyploids since *A. thaliana* is diploid while *B. rapa* is a recent allohexaploid (Wang et al., 2011). A common result of allopolyploidization is genome dominance, where the non-dominant subgenome experiences reduced expression and increased fractionation (Grover et al., 2012; Garsmeur et al., 2014). In *B. rapa*, the homeolog with more transposons within 500 bp of the transcription start or stop site was more likely to have lower expression and be present in the non-dominant subgenome (Woodhouse et al., 2014), suggesting that RdDM is associated with reduced expression from the non-dominant genome. This idea is supported by the higher proportion of transposon-derived 24-nt siRNAs mapping to regions surrounding genes of the non-dominant genome in *B. rapa*, cotton, and maize (Woodhouse et al., 2014; Renny-Byfield et al., 2015; Cheng et al., 2016; Forestan et al., 2017). Consequently, RdDM might be required to maintain the balance among subgenomes during reproduction, and increased expression of genes on the non-dominant subgenome in RdDM mutants might cause dosage imbalances that result in detrimental phenotypes (Alger and Edger, 2020). Species without a recent history of genome duplication have fewer homeologous gene pairs and less need for subgenome balance and thus might have fewer phenotypes when RdDM is eliminated.

Alternatively, RdDM’s role during reproduction might relate to balancing parental genomes in the endosperm, a process linked to RdDM (reviewed in (Chow and Mosher, 2023)). The triploid endosperm develops following the fertilization of the diploid central cell by a haploid sperm cell and therefore has a 2:1 maternal:paternal genome ratio. Distortion of this ratio results in failure during endosperm development, likely due to an imbalance in parental conflict over resource allocation to seeds (Scott et al., 1998; Haig, 2013). Similar imbalances are seen in interspecific crosses of the same ploidy, leading to the conclusion that it is not the *absolute* ploidy from each parent that determines balance, but rather the *effective* ploidy – a combination of ploidy and the strength of parental conflict (Johnston et al., 1980). The WISO hypothesis (Weak Inbreeder, Strong Outbreeder) proposes that parental conflict is proportional to the extent of outbreeding in a species (Brandvain and Haig, 2005; Brandvain and Haig, 2018). Until recent domestication, *B. rapa* is an outbreeding species, implying it will have strong conflict, while *A. thaliana* evolved inbreeding ∼ 500 thousand years ago (KYA) and should have weak conflict (de la Chaux et al., 2012). RdDM may be a necessary referee to mediate conflict between strong maternal and paternal interests in outbreeders, but loss of that balance is less important in inbreeders who have only weak conflict.

To test these hypotheses, we used a CRISPR/Cas9 DNA editing system to introduce mutations into two essential genes in the RdDM pathway in three Brassicaceae species that differed from each other in ploidy or breeding system. Our target species were: 1) *Camelina sativa*, a self-compatible recent allopolyploid; 2) the diploid obligate outcrosser *Capsella grandiflora*; and 3) *Capsella rubella*, a diploid inbreeder (**Supplementary Figure 1**). *C. sativa* underwent a polyploidization event ∼ 5.5 million years ago (MYA), retains three distinct subgenomes, and has a clear subgenomic expression bias (Kagale et al., 2014; Brock et al., 2018; Mandáková et al., 2019). We reasoned that if RdDM is critical in mediating conflict among subgenomes, *C. sativa* RdDM mutants would have severely reduced seed set similar to *B. rapa. C. rubella* split from its congener, *C. grandiflora*, < 200-50 KYA, soon after its founders lost self-incompatibility (Foxe et al., 2009; Guo et al., 2009; Slotte et al., 2013) yielding a fully self-compatible *C. rubella*. The sister relationship between *C. grandiflora* and *C. rubella* allowed us to test whether outcrossers exhibit a stronger requirement for RdDM during reproduction than do inbreeders. Here, we show that RdDM is not critical for seed production in *C. sativa*, suggesting that a recent history of polyploidy is not an important factor for RdDM mutant phenotypes. Instead, we observed a significant and striking reduction in seed set in *C. grandiflora*, suggesting that breeding system is a major factor determining reproductive impacts upon loss of RdDM.

## Results

### Creation of RdDM mutants in Brassicaceae species

To explore the role of RdDM during reproduction, we used CRISPR/Cas9-nickase (Fauser et al., 2014) to generate loss of function mutations in *RDR2* and *NRPE1* in three Brassicaceae species that vary in breeding system and history of genome duplication. *NRPE1* encodes the largest subunit of RNA Pol V (Ream et al., 2009). *C. rubella* and *C. grandiflora* encode single copies of *RDR2* and *NRPE1*, while *C. sativa* retains three copies of these genes due to its recent whole genome triplication. We identified guide RNAs that flank functionally important regions of the proteins and target all five versions of each gene (**Figure 1, Supplementary Figure 2**). Following floral-dip transformation of the genome editing constructs, T1 individuals were genotyped to identify strong mutations in target genes and these mutations were bred to homozygosity (**Supplementary Table 1, Supplementary Figure 2**). For *C. sativa*, mutations in all three gene copies were combined and the resulting triple mutants are hereafter designated *Cs rdr2-1, Cs rdr2-2, Cs nrpe1-1*, and *Cs nrpe1-2*. Similar to *rdr2* and *nrpe1* mutations in *A. thaliana* and *B. rapa*, the *C. sativa* and *C. grandiflora* mutants did not exhibit consistent vegetative growth defects, however all RdDM mutants in *C. rubella* were moderately smaller than wild type (**Figure 1**) (Grover et al., 2018).

**Figure 1:**
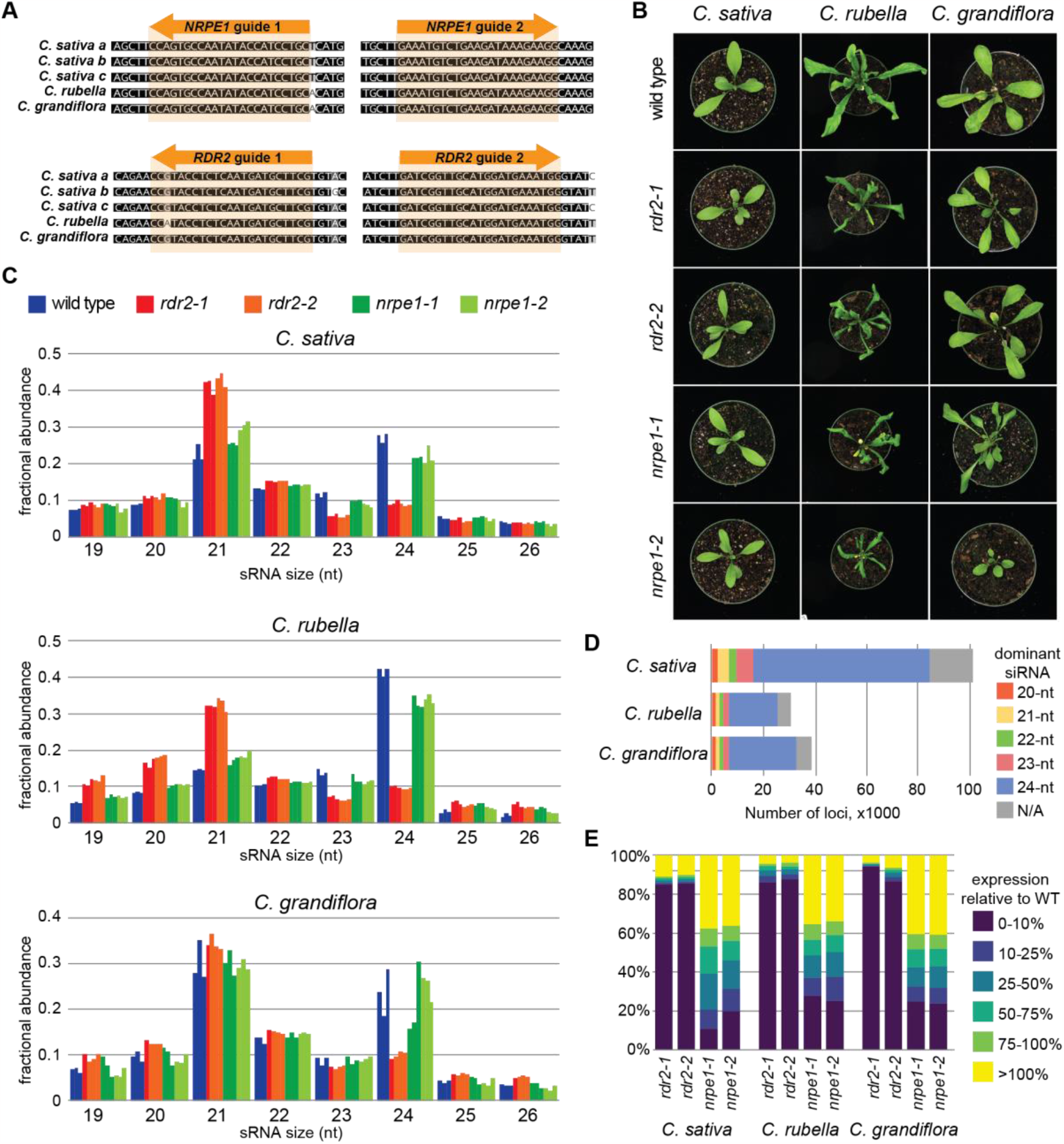
Creation and characterization of RdDM mutants in Brassicaceae species. (**A**) Alignment of guide RNA binding sites from *Camelina* and *Capsella NRPE1* and *RDR2* genes. (**B**) *Camelina* and *Capsella* RdDM mutants have no or mild vegetative growth phenotypes. (**C**) Size profiles of genome-mapping small RNAs from wild-type and RdDM mutant leaves of each species. Three biological replicates are shown for each allele, with the exception of *C. grandiflora rdr2-1*, which has only a single replicate. (**D**) The largest class of leaf siRNA loci in each species are 24-nt dominant loci. (**E**) In each species, *rdr2* mutations dramatically reduce expression from 80-90% of siRNA loci, while *nrpe1* mutations have a less pronounced effect on siRNA expression.

To confirm whether these mutations impact RdDM, we sequenced small RNAs from leaf tissue of each allele and mapped them to their respective genomes (**Supplementary Table 2**). To increase the proportion of mapped *C. grandiflora* sRNA reads, we used Oxford Nanopore Technology to sequence the genome of *C. grandiflora* accession 83.17, and then *de novo* assembled the reads, polished the assembly with publicly-available Illumina reads (Williamson et al., 2014), and annotated it. Our resequencing efforts yielded a *C. grandiflora* genome that is less fragmented compared to the previously published genome, with an increase in linearity to L/N50 of 17/2.3 Mb from 256/29.1 kb (**Supplementary Table 3**). The size profile of mapped sRNA reads demonstrates that there is strong loss of 23-24-nt siRNA in the *rdr2* lines of all three species, and a smaller reduction in 23-24-nt accumulation in *nrpe1* plants (**Figure 1**). A partial loss of siRNA accumulation in *nrpe1* is expected, as Pol V is not required for siRNA biosynthesis, but the failure to target methylation in *nrpe1* mutants results in reduced production of siRNAs via Pol IV (Kanno et al., 2005; Pontier et al., 2005; Mosher et al., 2008; Gouil and Baulcombe, 2016; Grover et al., 2018; Zheng et al., 2021). There is also an observable increase in all other size classes in the *rdr2* mutants, likely due to over-sampling these species following loss of the major size class.

Next, we used ShortStack to define small (s)RNA-producing loci in each genome. The *Capsella* species have 30-40,000 sRNA loci, while *C. sativa* has approximately 3 times that number, perhaps due to its recent whole genome triplication (**Figure 1**). Categorizing these loci by the dominant size of sRNA demonstrates that 24-nt sRNA loci form the largest group (61-68%). We then assessed the accumulation of sRNA from each locus in the mutants. In each species, *rdr2* mutant exhibit a strong reduction in sRNA from 80-90% of sRNA loci and increased sRNA accumulation from a small proportion of loci (**Figure 1**). In contrast, fewer sRNA loci were strongly reduced in *nrpe1* mutants, and many loci displayed only a moderate reduction in sRNA accumulation. This pattern is consistent with sRNA size profiles and sRNA locus analysis of similar mutations in *A. thaliana*, tomato, *B. rapa*, and rice (Mosher et al., 2008; Gouil and Baulcombe, 2016; Grover et al., 2018; Zheng et al., 2021), indicating that the mutations generated here disrupt the RdDM pathway.

### Seed development is disrupted upon loss of RdDM in outbreeding *C. grandiflora*

To investigate reproductive phenotypes upon loss of RdDM, we first assessed seed set, defined here as the number of healthy seeds per fruit. Because *C. sativa* fruits do not lose their valves (shatter) at maturity, we counted the number of healthy seeds in mature fruits. Seeds that were shriveled or darkly pigmented were considered inviable (**Supplementary Figure 3**). Fruits in both *Capsella* species shatter, particularly when they contain many mature seeds. For these species we therefore counted developing seeds at 7 days after pollination (7 DAP), a timepoint that allows us to measure seed development before shattering. Seeds in these fruits were considered healthy if they were plump and green; white, brown, and/or unexpanded seeds were counted as aborted or unfertilized (**Supplementary Figure 3**).

RdDM mutant lines in all three species exhibited reduced seed set (**Figure 2**), however the scale of reduction varied. *Cs rdr2* plants had 25-32% reduction in seed set compared with wild type, while *Cs nrpe1-1* showed no change in seed set. RdDM mutations in *C. rubella* reduced healthy seeds at 7 DAP by approximately half. In comparison, *C. grandiflora rdr2* alleles had 97-99% fewer healthy seeds in the RdDM mutant fruits compared to wild type, while *Cg nrpe1* mutants showed approximately 70% reduction. We also measured the weight of mature seeds collected in bulk, which demonstrated that even seeds scored as healthy are slightly smaller for RdDM mutants (**Figure 2**). Nevertheless, mature seeds of each mutant routinely germinated at high frequency. Taken together with observations in *A. thaliana* and *B. rapa*, these results indicate that the RdDM pathway plays a role during seed development in each of the species tested, with the strongest effect in outbreeding *C. grandiflora*.

**Figure 2.**
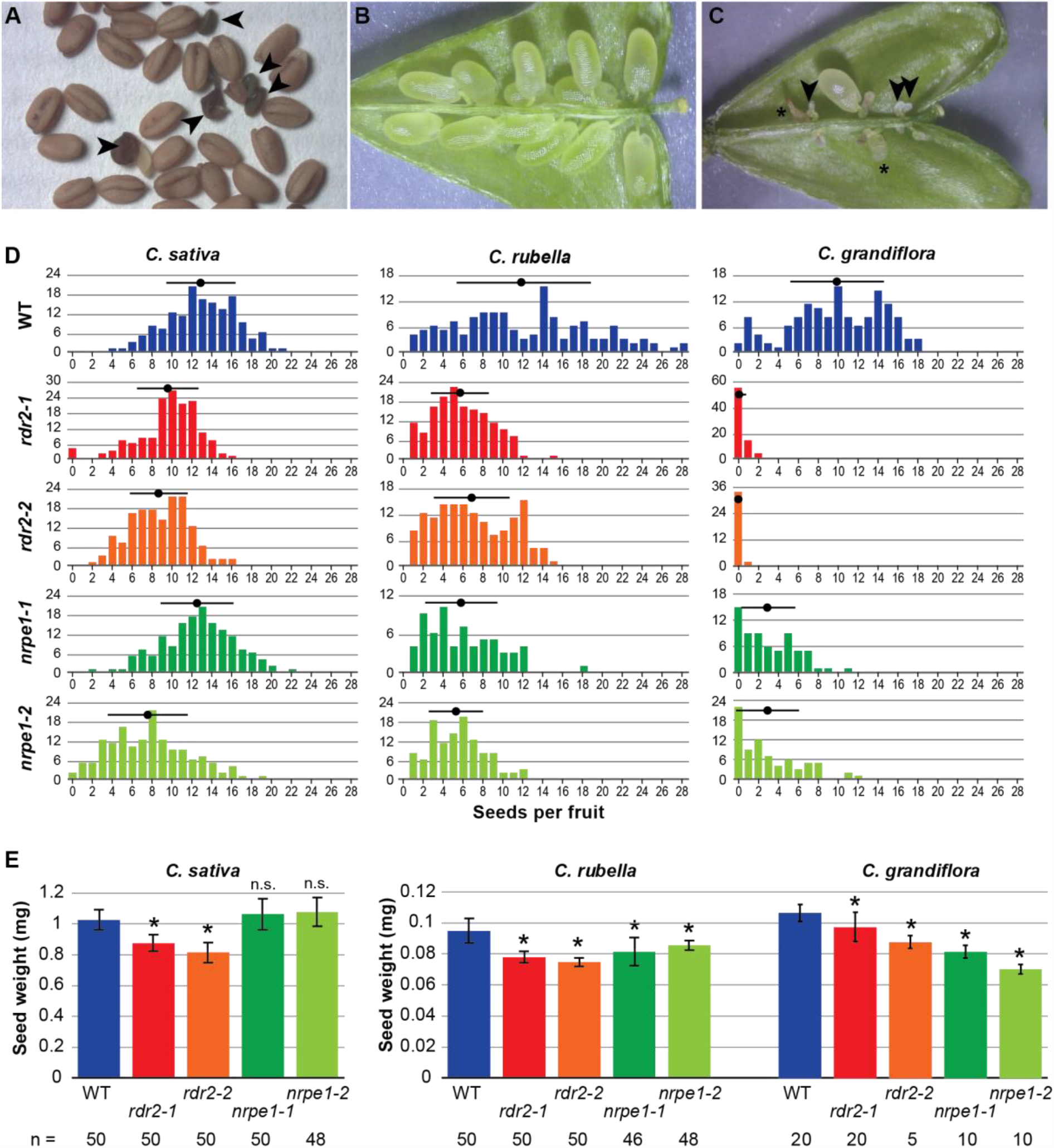
Seed production in Brassicaceae RdDM mutants. (**A**) Example seeds from *C. sativa*. Arrowheads indicate seeds scored as unhealthy. (**B, C**) Example seeds in *C. grandiflora* wild type and RdDM mutant fruits at 7 DAP. Arrowheads point to unfertilized ovules and asterisks mark aborted seeds. (**D**) Histograms of the number of mature seeds per fruit in *Camelina* or healthy seeds at 7 DAP in *Capsella*. The average seed count is shown with a black dot; black bars represent the standard deviation around the mean. (**E**) Seed weight of mature seeds. Pools of 20 seeds were weighed three times and the average of each pool was divided by the number of seeds to obtain an average seed weight. The average and standard deviation of these seed weights for n pools are shown. ^*^ = p<0.01 after Bonferroni FDR correction; n.s. = not significant.

### Maternal factors are responsible for seed set reductions in *C. rubella* and *C. grandiflora*

To better understand the role of RdDM in species with different mating systems, we further compared the reproductive phenotypes of RdDM mutants in the two *Capsella* species. Wang et al. (2020) reported defective pollen in *C. rubella nrpd1* mutants, with approximately 80% of the pollen arresting prior to maturity (Wang et al., 2020). We therefore imaged mature pollen grains from each *C. rubella* and *C. grandiflora* RdDM mutant line to determine if reduction in seed set resulted from defects in pollen maturation. We observed no difference in the proportion of mature pollen compared to wild type (**Figure 3A**). We also tested pollen viability with fluorescein diacetate staining and observed no difference in viability between wild type and mutants (**Figure 3**). Together these observations suggest that defects in pollen development were not responsible for the observed decrease in seed set for any *Capsella* RdDM mutant. Moreover, pollination with wild-type pollen was unable to rescue the seed production defect in any of the *Capsella* RdDM mutant lines (**Figure 4**). These observations indicate that decreased seed production in *Capsella* RdDM mutants is primarily due to either a maternal sporophytic or female gametophytic defect, a feature that is shared with seed production phenotypes of *B. rapa* RdDM mutants (Grover et al., 2018).

**Figure 3:**
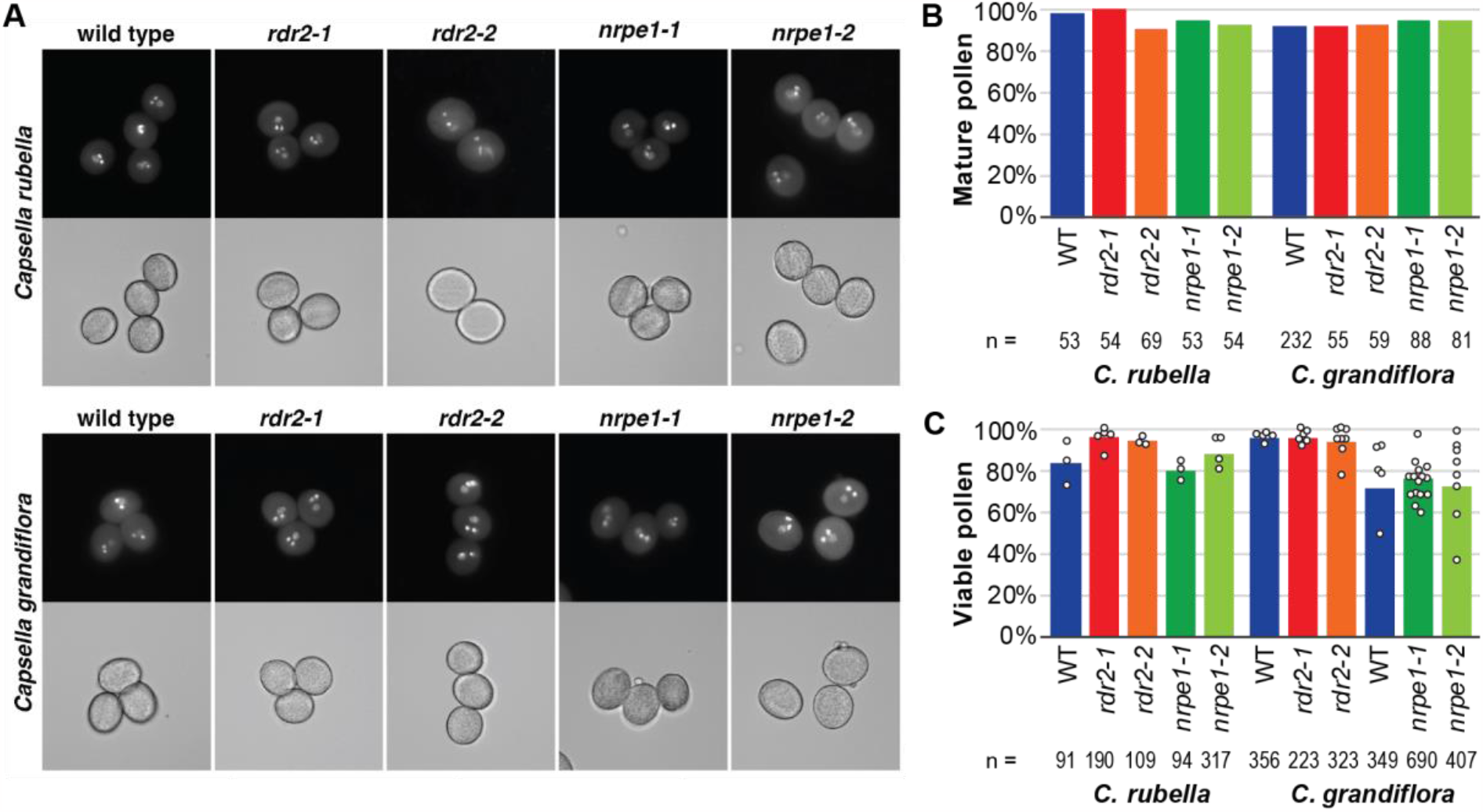
Normal pollen development in *Capsella* RdDM mutants. (**A**) Representative images of pollen from *Capsella* RdDM mutants demonstrates that most pollen is mature and trinucleate, with a single diffuse vegetative nucleus and two condensed sperm cells (top, DAPI; bottom, brightfield). (**B**) Quantification of mature (trinucleate) pollen and the number of assessed pollen grains. (**C**) Quantification of viable pollen by FDA staining for all tested pollen grains, with the total number of grains listed below. Circles represent measurements from individual slides or groups of slides. *C. grandiflora nrpe1* and *rdr2* were collected separately, and therefore each is compared to a synchronous wild type control.

**Figure 4:**
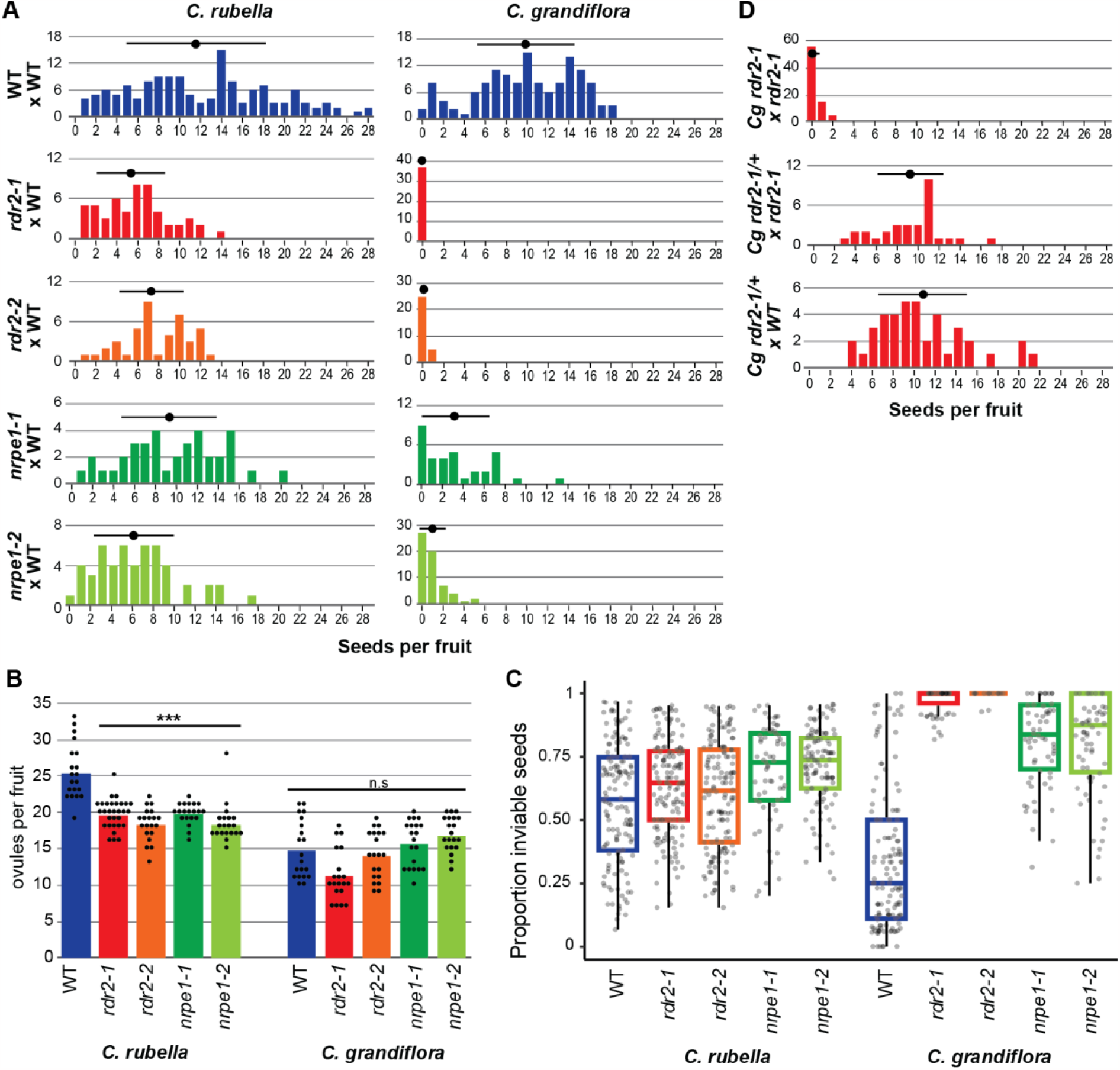
Seed set reduction in *Capsella* RdDM mutants is controlled maternally. (**A**) Number of “healthy” seeds per fruit at 7 DAP in *Capsella* RdDM mutants crossed with wild-type pollen. WT x WT are the same data presented in Figure 2A. (**B**) Number of ovules in unfertilized pistils (n > 20). Colored bars represent the average; black dots are individual datapoints. All *C. rubella* RdDM mutants are significantly different from wild type (WT) at p < 0.01 after Bonferroni correction (Student’s T test). *C. grandiflora* mutants are not significantly different at this threshold. (C) Fraction of things inside a fruit at 7 DAP that are not scored as “healthy”, representing either failure to fertilize or seed abortion. (**D**) Number of “healthy” seeds per fruit at 7 DAP in *C. grandiflora rdr2* heterozygotes crossed with homozygous mutant or wild-type pollen. *Cg rdr2-1* x *Cg rdr2-1* are the same data presented in Figure 2A.

Reduction in seed set might result from fewer ovules, decreased fertilization, or increased seed abortion. To distinguish among these possibilities, we analyzed the number of ovules in unfertilized pistils from each mutant (**Figure 4**). In *C. grandiflora*, there is no statistically significant difference in ovule number between wild type and RdDM mutants, however *C. rubella[para-merge-che* RdDM mutants exhibit approximately 25% fewer ovules than wild type (p < 0.01, Bonferonni corrected Student’s T test), a reduction that accounts for most of the reduced seed set in these mutants. The higher number of ovules in *C. rubella* wild type relative to *C. grandiflora* wild type is consistent with the decrease in pollen:ovule ratio commonly observed following the transition to self-fertilization (Sicard and Lenhard, 2011). To further explore the seed set reduction in *Capsella* RdDM mutants, we counted the number of inviable seeds in 7 DAP fruits and expressed this as a fraction of all objects in a fruit to account for differences in ovule number (**Figure 4**). Inviable seeds included those that had clearly begun development before aborting as well as small white seeds that might be unfertilized ovules or result from abortion shortly after fertilization (**Figure 2**). While *C. rubella* plants displayed relatively high levels of seed failure, this was independent of RdDM genotype. Despite this fact, we cannot entirely rule out a role for gametophytic incompatibilities transmitted from *C. rubella* pollen during fertilization, though our data indicate that the effect of such incompatibilities is minor relative to the effect of maternal ovule production. In contrast, *C. grandiflora rdr2* and *nrpe1* fruits exhibited substantially higher levels of seed failure than wild type (**Figure 4**), including 14-56% clearly resulting from seed abortion. From this we conclude that reduction in seed set in *C. rubella* RdDM mutants is primarily due to the production of fewer ovules before fertilization, while abortion after fertilization is the major contributor to reduced seed set observed in *C. grandiflora* mutants.

*B. rapa* RdDM mutants also exhibit extensive seed abortion after fertilization, a trait that is controlled by the maternal sporophytic genotype (Grover et al., 2018). To test whether seed abortion is sporophytically determined in *C. grandiflora*, we crossed heterozygous plants with wild-type or homozygous mutant pollen and scored seed development at 7 DAP. Seed production in such crosses is indistinguishable from wild type (**Figure 4**), indicating that mutant embryo sacs develop normally even when fertilized by mutant pollen, as long as they are supported by a maternal sporophyte capable of RdDM.

## Discussion

We sought to test two hypotheses to explain the different seed production phenotypes of RdDM mutants in Brassicaceae. First, that failure of seed development in RdDM mutants was associated with recent polyploidy and ongoing balance between subgenomes, and secondly, that breeding system and the increased parental conflict expected in outbreeding species results in a requirement for RdDM. Here, we show that RdDM is critical for seed development in the outbreeding *C. grandiflora* while its role is less important in inbreeding species, including inbreeding polyploids.

Seed production was disrupted when RdDM was eliminated in the three Brassicaceae species, but to a different extent in each (**Figure 2**). RdDM mutants in polyploid *C. sativa* had only a very slight decrease in seed set. RdDM might be important for maintaining balance among its three subgenomes, however loss of this balance does not confer a reproductive phenotype. Similarly, although loss of RdDM causes a reduced size of rosette and fewer ovules in the diploid inbreeder *C. rubella* (**Figure 1, Figure 4**), upon fertilization, those ovules produce healthy seeds at the same frequency as wild type plants (**Figure 4**). In contrast, *C. grandiflora* RdDM mutants had a strong post-fertilization defect that was dependent of the maternal sporophytic genotype; this defect showed similar characteristics and genetic control as RdDM mutants in the recently outbreeding *B. rapa* (Grover et al., 2018).

A maternal sporophytic defect impacting seed number and size in *C. rubella nrpd1* mutants has been reported (Wang et al., 2020), however it is not clear whether this results from a decreased number of ovules (as we detect for *Cr rdr2* and *Cr nrpe1*) or another mechanism. Wang et al. (2020) also observed a reduction in the proportion of viable pollen in *C. rubella nrpd1* mutants. We do not detect a pollen defect in either *C. rubella* or *C. grandiflora* RdDM mutants (**Figure 3**). Combined with the fact that wild-type pollen is unable to rescue seed production from *Capsella* RdDM mutant mothers (**Figure 4**), this observation suggests that male gametophytic effects are unlikely to contribute to the RdDM mutant phenotypes we documented. This discrepancy in pollen phenotype for *C. rubella* RdDM mutants might be due to the mutation of different siRNA biosynthesis genes (*NRPD1* versus *RDR2*), however these proteins function together to generate dsRNA and would be expected to yield highly similar, if not identical, phenotypes (Singh et al., 2019; Fukudome et al., 2021; Huang et al., 2021). Variation in pollen phenotypes might also result from the use of different *C. rubella* accessions or different growth conditions.

The requirement for maternal sporophytic RdDM for successful seed production in *C. grandiflora* implicates maternally-derived siRNAs. The most abundant of these are siren siRNAs, which are expressed in the developing seed coat and might be trafficked into the developing endosperm (Grover et al., 2020). Siren siRNAs trigger non-CG methylation of protein-coding genes, which can result in altered gene expression (Burgess et al., 2022). Further research is needed to determine whether ovule siRNA populations and levels of methylation differ between *C. rubella* and *C. grandiflora*, or whether specific sRNA loci are required for seed development in outbreeding species.

Loss of RdDM has been linked to reduction of effective ploidy in diploid by tetraploid crosses in *A. thaliana* (Erdmann et al., 2017; Martinez et al., 2018; Satyaki and Gehring, 2019). Interspecific crosses indicate that *C. grandiflora* has a higher effective ploidy than *C. rubella*, which is consistent with the WISO hypothesis and indicative of stronger parental conflict in *C. grandiflora* (Brandvain and Haig, 2005; Rebernig et al., 2015; Brandvain and Haig, 2018; Lafon-Placette et al., 2018). We therefore hypothesize that loss of RdDM reduces effective ploidy and disrupts the balance between parental genomes after fertilization. For outbreeding species with high effective ploidy, this disruption results in a greater developmental defect because parental conflict is high. Inbreeding species have limited parental conflict and therefore disruption of the maternal:paternal ratio has less impact.

Transposable element content, and the proximity of transposons to protein-coding genes, has also been proposed to explain variation in phenotypes upon loss of RdDM in different species (Wei et al., 2014; Gouil and Baulcombe, 2016; Wang et al., 2020). However, *C. rubella* and *C. grandiflora* genomes have similar transposon number, expression, and proximity to genes (Slotte et al., 2013; Agren et al., 2014), and both *Capsella* genomes have a much lower transposon content than *B. rapa* (Wang et al., 2011). Therefore, transposon density cannot account for the dramatic difference in reproductive phenotype when RdDM is eliminated, although we cannot eliminate the possibility that it is linked to the likelihood of vegetative phenotypes.

There are a suite of floral development changes that commonly occur following the transition to self-compatibility, together referred to as the selfing syndrome (Sicard and Lenhard, 2011). Reduction in parental conflict, effective ploidy, and the requirement for RdDM could therefore be considered an epigenomic selfing syndrome – a suite of epigenomic changes that commonly develop over evolutionary time following the transition to self-fertilization. Our work provides the foundation for further study of this phenomenon.

## Methods

### Plant material and growth conditions

*C. sativa* cultivar Ames (a gift of John McKay, Colorado State University), *C. rubella* Monte Gargano, and *C. grandiflora* accession 83.17 (both donated by Stephen Wright, University of Toronto) were grown at 20°C under 16h:8h light:dark conditions. *C. sativa* seeds were stratified for one day at 4°C prior to germination; *C. rubella* and *C. grandiflora* were both stratified for two weeks at 4°C. *C. rubella* also required three weeks of vernalization at 4°C to induce flowering. For the collection of unfertilized ovules, *C. rubella* flowers were emasculated approximately 24 hours before anthesis; pistils were manually dissected 2 days later. *C. grandiflora* ovules were collected identically to *C. rubella* except emasculation was not performed.

### CRISPR design and plasmid construction

The T-DNA cassette used for transformation contained Cas9 nickase under a ribosomal protein promoter (*AtRPS5A::Cas9-D10A*), two guide RNAs, EGFP-tagged oleosin under its native promoter (*AtOLE::OLE-GFP*), and a BASTA resistance gene. The vector was constructed using traditional cloning techniques in *E. coli* to combine desired promoters, reporters, and proteins into a single vector with a variety of source materials. Assembled T-DNA plasmids were purified and transformed into *Agrobacterium tumefaciens* GV3101.

Two protospacers for both *RDR2* and *NRPE1* were selected manually from regions conserved in all three species with target specificity verified using CRISPOR (Haeussler et al., 2016) and CRISPRdirect (Naito et al., 2015). The protospacers for *RDR2* were 5’-ccrTACCTCTCAATGATGCTTCG - 3’ and 5’ - GATCGGTTGCATGGATGAAAtgg - 3’ (PAM sequences in lowercase). The protospacers for *NRPE1* were 5’ - ccaGTGCCAATATACCATCCTGC - 3’ and 5’ - GAAATGTCTGAAGATAAAGAagg - 3’.

### Plant transformation, genotyping, and phenotyping

*C. grandiflora* plants were transformed using *Agrobacterium*-mediated floral dip as described in (Dew-Budd et al., 2019). Briefly, developing fruits and opened flowers were removed prior to the first *Agrobacterium* dipping. Plants were dipped 3-4 times at an interval of 4-7 days, and were hand pollinated daily following the dipping. *C. rubella* and *C. sativa* were transformed via standard floral dip (Clough and Bent, 1998). Positive transformants were selected through EGFP visualization in mature seeds using a Zeiss Axiozoom 16 fluorescent stereo microscope with a 488 nm filter.

RdDM mutants were genotyped with PCR primers flanking the protospacers (**Supplementary Table 4**). Size differences between wild-type and mutant amplicons were visualized on an agarose gel and bands were sequenced to identify the exact mutation.

For *C. sativa*, dried fruits were collected from five individuals of each genotype to measure seed set. To prevent bias due to preferential dehiscence of the largest fruits, *C. rubella* and *C. grandiflora* seed set was measured 7 days after manual pollination (DAP), which is approximately torpedo stage in wild type. For *C. grandiflora* all plants were genotyped and only crossed if they shared no more than one S allele to avoid confounding effects from S-locus incompatibility. Upon maturation, seeds were dried for at least two weeks under vacuum before measuring in batches of twenty on an ultra-microbalance (Sartorius Cubis MSE3.6P-000-DM).

For DAPI staining, pollen grains were harvested by vortexing and centrifuging flowers in pollen isolation buffer (100 mM NaPO_4_, pH 7.5; 1 mM EDTA; 0.1% (v/v) Triton X-100. The flowers and the fixing solution were removed and replaced with 1 µg/mL DAPI (4′,6-diamidino-2-phenylindole, BD Biosciences) diluted in pollen isolation buffer before visualizing under a fluorescence microscope using a 420 nm filter (Zeiss LSM880 Inverted Confocal Microscope). Pollen was considered normal if two sperm cell nuclei were distinguishable from the vegetative nucleus. The majority of abnormal pollen were small and did not fluoresce. FDA staining for pollen viability was performed according to (Li, 2011).

### *Capsella grandiflora* genome sequencing, assembly, and annotation

Genomic DNA was isolated using a modified protocol of the Circulomics Nanobind Plant Nuclei Big DNA Kit to account for *C. grandiflora*’s small genome. The modified extraction replaced the nuclei isolation with Circulomics’ supplementary direct tissue lysis protocol. Approximately 1 µg of genomic DNA was prepared using the Oxford Nanopore LSK-109 kit without shearing and sequenced on a MinION system using a R9.4 flow cell. Reads were called during sequencing using Guppy (v3.2.6) with a MinIT device (ont-minit-release 19.10.3) used for processing.

Adaptors were trimmed from the reads with Porechop (v0.2.3, https://github.com/rrwick/Porechop) using default parameters, and then filtered for reads > 500 bp in length with an average quality > 9 using Nanofilt (v2.6.0) (De Coster et al., 2018). Reads matching either the *C. grandiflora* chloroplast genome or the *C. rubella* mitochondrial genome were removed using minimap2 (v2.17) (Li, 2018) and SAMtools (v1.4) (Li et al., 2009). The remaining reads were *de novo* assembled using Canu (v1.9) (Koren et al., 2017) with default parameters and an estimated genome size of 120 Mb. The resulting genome was polished using the filtered long read data with Racon (v.1.4.3, non-default parameters: -m 8 -x 6 -g 8) (Vaser et al., 2017) and Medaka (v0.12.1, https://github.com/nanoporetech/medaka). Accession-specific publicly available short-read data from Williamson (2014, SRR1508428) was used to reiteratively polish three times with Pilon (v1.23) (Walker et al., 2014; Williamson et al., 2014). Due to the heterozygosity of *C. grandiflora*, the polished genome was further processed with Purge Haplotigs to separate the main genome and the haplotigs (Roach et al., 2018). Thirty-seven contigs less than 1 kb were removed from the main assembly and seven contigs less than 1 kb were removed from the haplotig assembly. The assembly is available at NCBI (BioProject PRJNA988139).

### Small RNA-seq generation and analysis

Small RNA sequencing libraries were prepared using a standard protocol (Grover et al., 2018). Briefly, after RNA precipitation, small RNA was enriched with a mirVana miRNA isolation kit (Thermo Fisher Scientific) followed by NEBNext small RNA library preparation (New England Biolabs). Three independent biological replicates were prepared for each genotype then sequenced at the University of Arizona Genomics Core on a NextSeq500. Raw reads are available at NCBI SRA (BioProject number PRJNA986946).

Small RNA reads were processed and aligned to the appropriate genome using the sRNA snakemake workflow (v1.0) (Bose and Grover, 2019). Briefly, raw reads were trimmed for size and quality using Trim Galore! (v0.6.2) (Krueger, 2012), contaminating non-coding RNAs were removed using bowtie (v1.2.2) (Langmead et al., 2009) with the Rfam database entries for *A. thaliana* and *C. rubella* (Kalvari et al., 2018), then reads aligning to the species-specific chloroplast and *C. rubella* mitochondrial genomes were removed with bowtie. The remaining reads were aligned to the reference genome using ShortStack (v3.8.5) (Axtell, 2013) with the snakemake workflow’s default parameters except one mismatch was allowed during ShortStack alignment. Biological replicates were pooled for locus-level analyses.

## Acknowledgements and Funding

The authors wish to thank the following colleagues for their assistance and expertise: University of Arizona Genomics Core for Illumina sequencing, Professor David Baltrus (University of Arizona) for Nanopore expertise, Professor Steven Wright (University of Toronto) for seeds of *Capsella* species, Professor John McKay (Colorado State University) for *Camelina sativa* seed, Arizona Laboratory for Emerging Contaminants for access to the microbalance and Danielle Barrientes for training on its use, and undergraduate assistants Jeff Clark, Jack Stearns, and Cecelia Martinez. The authors are grateful for support from the National Science Foundation (IOS-1546825 to R.A.M. and M.A.B.), the USDA National Institute of Food and Agriculture (AFRI 2021-67013-33797 to R.A.M.), the USDA Hatch funding (ARZT 1361510-H25-249 to R.A.M and M.A.B), and The University of Arizona College of Agriculture and Life Sciences iVIP award (to M.A.B.).

## Author Contributions

RAM and MAB designed the research; KDB, HTC, TK, BCD, and JAR performed research; KDB, HTC, RAM, and MAB analyzed data and wrote the manuscript.

