## Supplementary for "Mating system is associated with seed phenotypes upon loss of RNA-directed DNA Methylation in Brassicaceae"

**Supplementary Table 1: Alleles created in this study**

| Gene | Gene_ID[1] | Mutation | First altered residue | Effect of mutation[2] |
|---|---|---|---|---|
| ***Cg nrpe1-1*** | Cagra.2495s0018.1 | 12 bp deletion | K 100 I | Deletes Zn binding domain |
| ***Cg nrpe1-2*** | Cagra.2495s0018.1 | 51 bp deletion | K 100 N | Deletes Zn binding domain and splice site |
| ***Cr nrpe1-1*** | LOC17889800 | 16 bp deletion | I 81 M | Deletes Zn binding domain + Frameshift |
| ***Cr nrpe1-2*** | LOC17889800 | 46 bp insertion | C 98 P | Deletes Zn binding domains + Frameshift |
| ***Cs nrpe1-1*** | LOC104699339 | 43 bp insertion | C 98 S | Deletes Zn binding domains + Frameshift |
| | LOC104788861 | 34 bp insertion | C 101 T | Deletes Zn binding domain + Frameshift |
| | LOC104785486 | 29 bp insertion | L 102 O | Frameshift in RPB domain 1 |
| ***Cs nrpe1-2*** | LOC104699339 | 37 bp deletion | M 93 R | Deletes Zn binding domains + Frameshift |
| | LOC104788861 | 65 bp insertion | K 103 Y | Frameshift in RPB domain 1 |
| | LOC104785486 | 49 bp insertion | L 102 S | Frameshift in RPB domain 1 |
| ***Cg rdr2-1*** | Cagra.16778s0004.1 | 31 bp deletion | G 729 W | Frameshift |
| ***Cg rdr2-2*** | Cagra.16778s0004.1 | 18 bp deletion | L 703 V | Deletes most of helix 11 |
| ***Cr rdr2-1*** | LOC17882313 | 89 bp insertion | C 735 Stop | Frameshift |
| ***Cr rdr2-2*** | LOC17882313 | 44 bp insertion | E 716 D | Frameshift |
| ***Cs rdr2-1*** | LOC104719267 | 174 bp insertion | C 737 M | Triplicates helix 12, Strand 12 - 13 |
| | LOC104707406 | 90 bp insertion | C 737 M | Duplicates helix 12, Strand 12-13 |
| | LOC104737110 | 22 bp deletion | P 700 V | Frameshift |
| ***Cs rdr2-2*** | LOC104719267 | 180 bp insertion | G 736 S | Triplicates helix 12, Strand 12 - 13 |
| | LOC104707406 | 115 bp insertion | C 737 M | Insertion with a Stop codon |
| | LOC104737110 | 40 bp deletion | L 724 K | Frameshift |

[1] Gene IDs for *C. rubella* and *C. sativa* are from NCBI; gene IDs for *C. grandiflora* are from Phytozome

[2] Domain and structural information were ascertained from Cramer, et al. 2001 and Lakshminarayan, et al. 2003 for NRPE1 and RDR2, respectively

### Supplementary Table 2: sRNA sequencing stats

| Species | Genotype | Raw reads | structural RNAs removed | 19-26-nt filtered reads | Mapped 19-26-nt filtered reads |
|---|---|---|---|---|---|
| ***C. sativa*** | WT_A_leaf | 18,594,320 | 6,655,384 | 5,328,452 | 3,119,544 |
| ***C. sativa*** | WT_B_leaf | 21,259,632 | 5,062,896 | 3,698,912 | 2,116,148 |
| ***C. sativa*** | WT_C_leaf | 23,171,824 | 7,085,408 | 4,819,716 | 2,697,988 |
| ***C. sativa*** | rdr2-1_A_leaf | 22,477,136 | 3,607,804 | 1,151,136 | 601,600 |
| ***C. sativa*** | rdr2-1_B_leaf | 22,044,896 | 3,568,888 | 1,410,176 | 779,552 |
| ***C. sativa*** | rdr2-1_C_leaf | 28,748,644 | 5,444,752 | 1,713,552 | 958,392 |
| ***C. sativa*** | rdr2-2_A_leaf | 19,328,196 | 2,599,120 | 1,291,372 | 730,532 |
| ***C. sativa*** | rdr2-2_B_leaf | 22,851,068 | 2,687,068 | 1,457,288 | 840,380 |
| ***C. sativa*** | rdr2-2_C_leaf | 25,749,896 | 3,446,240 | 1,380,268 | 810,056 |
| ***C. sativa*** | nrpe1-1_A_leaf | 22,718,556 | 5,997,352 | 3,856,968 | 2,146,488 |
| ***C. sativa*** | nrpe1-1_B_leaf | 22,460,100 | 3,644,432 | 1,914,904 | 1,071,828 |
| ***C. sativa*** | nrpe1-1_C_leaf | 16,495,796 | 3,618,620 | 2,425,524 | 1,196,732 |
| ***C. sativa*** | nrpe1-2_A_leaf | 24,100,472 | 5,242,216 | 3,122,536 | 1,885,668 |
| ***C. sativa*** | nrpe1-2_B_leaf | 27,719,544 | 6,648,684 | 3,417,116 | 1,942,396 |
| ***C. sativa*** | nrpe1-2_C_leaf | 21,389,684 | 4,979,196 | 3,702,016 | 2,034,060 |
| ***C. rubella*** | WT_A_leaf | 11,367,024 | 2,434,844 | 1,247,828 | 587,544 |
| ***C. rubella*** | WT_B_leaf | 19,043,748 | 3,539,804 | 1,754,968 | 788,100 |
| ***C. rubella*** | WT_C_leaf | 18,636,148 | 2,164,348 | 1,517,760 | 747,580 |
| ***C. rubella*** | rdr2-1_A_leaf | 24,475,984 | 4,037,304 | 1,196,296 | 902,280 |
| ***C. rubella*** | rdr2-1_B_leaf | 19,507,360 | 2,729,360 | 1,243,844 | 608,904 |
| ***C. rubella*** | rdr2-1_C_leaf | 19,860,696 | 2,682,028 | 1,715,832 | 832,540 |
| ***C. rubella*** | rdr2-2_A_leaf | 17,933,924 | 2,693,980 | 1,298,052 | 681,820 |
| ***C. rubella*** | rdr2-2_B_leaf | 23,244,164 | 2,557,552 | 1,195,932 | 615,812 |
| ***C. rubella*** | rdr2-2_C_leaf | 18,187,316 | 1,807,464 | 1,195,932 | 608,796 |
| ***C. rubella*** | nrpe1-1_A_leaf | 20,772,948 | 2,363,788 | 1,573,064 | 722,028 |
| ***C. rubella*** | nrpe1-1_B_leaf | 19,333,952 | 3,144,284 | 1,962,316 | 871,676 |
| ***C. rubella*** | nrpe1-1_C_leaf | 20,799,540 | 3,524,592 | 1,824,876 | 787,256 |
| ***C. rubella*** | nrpe1-2_A_leaf | 26,491,300 | 4,885,372 | 2,843,096 | 1,388,436 |
| ***C. rubella*** | nrpe1-2_B_leaf | 22,343,912 | 4,740,344 | 2,815,160 | 1,416,128 |
| ***C. rubella*** | nrpe1-2_C_leaf | 17,850,148 | 3,097,904 | 2,245,720 | 1,030,408 |
| ***C. grandiflora*** | WT_A_leaf | 20,893,860 | 3,762,276 | 1,909,336 | 847,884 |
| ***C. grandiflora*** | WT_B_leaf | 18,939,788 | 2,865,772 | 1,615,708 | 777,280 |
| ***C. grandiflora*** | WT_C_leaf | 19,452,536 | 5,431,368 | 2,596,076 | 1,146,296 |
| ***C. grandiflora*** | rdr2-1_A_leaf | 19,305,768 | 2,274,588 | 874,736 | 353,064 |
| ***C. grandiflora*** | rdr2-2_A_leaf | 20,308,076 | 2,872,732 | 1,190,048 | 538,276 |
| ***C. grandiflora*** | rdr2-2_B_leaf | 20,718,180 | 3,360,744 | 1,091,232 | 524,036 |
| ***C. grandiflora*** | rdr2-2_C_leaf | 18,407,368 | 1,993,192 | 806,936 | 285,360 |
| ***C. grandiflora*** | nrpe1-1_A_leaf | 17,708,280 | 4,730,840 | 3,212,200 | 1,608,896 |
| ***C. grandiflora*** | nrpe1-1_B_leaf | 22,667,316 | 2,643,240 | 1,373,192 | 601,416 |
| ***C. grandiflora*** | nrpe1-1_C_leaf | 19,740,952 | 3,568,852 | 1,834,512 | 876,092 |
| ***C. grandiflora*** | nrpe1-2_A_leaf | 20,665,208 | 5,132,752 | 3,022,660 | 1,457,988 |

| | | | | | |
|---|---|---|---|---|---|
| ***C. grandiflora*** | nrpe1-2_B_leaf | 16,649,116 | 4,986,680 | 2,818,276 | 1,396,064 |
| ***C. grandiflora*** | nrpe1-2_C_leaf | 19,481,060 | 4,580,936 | 1,944,676 | 888,544 |

### Supplementary Table 3: *C. grandiflora* genome stats

| | Species: | *C. rubella* | *C. grandiflora* | *C. grandiflora* |
|---|---|---|---|---|
| | Source: | Slotte, et al 2013 (2) | Slotte, et al 2013 | This publication |
| Scaffolds | Total | 773 | 30,490 | 577 |
| | L50 | 4 | 256 | 17 |
| | N50 | 15.1 Mb | 98.1 Kb | 2.3 Mb |
| BUSCO Scores (1) | % Complete | 98.5% | 94.6% | 97.8% |
| | % Fragmented | 0.1% | 1.3% | 0.4% |
| sRNA reads | Mapped uniquely | 45.2% | 26.5% | 36.7% |
| | Multi-mapped | 27.5% | 7.3% | 26.3% |

(1) Compared to Brassicales BUSCOs
(2) NCBI Assembly GCA_000375325.1

### Supplementary Table 4: Genotyping primers

| Gene | Species | Forward | Reverse |
|---|---|---|---|
| **RDR2** | *C. grandiflora* | TCCGTCGAGATCACCACCA | TATGTCCCTTCTGCATTTCAAATTCG |
| | *C. rubella* | TCCGTCGAGATCACCACCA | TATGTCCCTTCTGCATTTCAAATTCG |
| | *C. sativa* (A) | AAGTGTTTGAGGCGATGCAAG | CAACAAAAACCAAGGAGTCTTAG |
| | *C. sativa* (B) | CAGTGTTTGAGGCCATGCAAG | ACCAAAGTAGAAAGAAGTTATCAAC |
| | *C. sativa* (C) | AAGTGTTTGAGGCGATGCAAG | GCATAGCTTTAAAGGATTGATTTAACATATA |
| **NRPE1** | *C. grandiflora* | TCAACCATGCCAGTCAACTCTCTA | CAACTTTATATGGCATTACTCACCTCA |
| | *C. rubella* | TCAACCATGCCAGTCAACTCTCTA | CAACTTTATATGGCATTACTCACCTCA |
| | *C. sativa* (A) | TCTTTCTACCTACCCTCCAAG | AAGCACAATTACTATATGCAACTAG |
| | *C. sativa* (B) | GTTTTGCAATTTTTTGGTTCCA | AAGCACAATTACTATATGCAACTAG |
| | *C. sativa* (C) | GGTTATGCTTGCCACTAAATTC | ATAGATATGAATTAGCACGTTTTCTAAC |

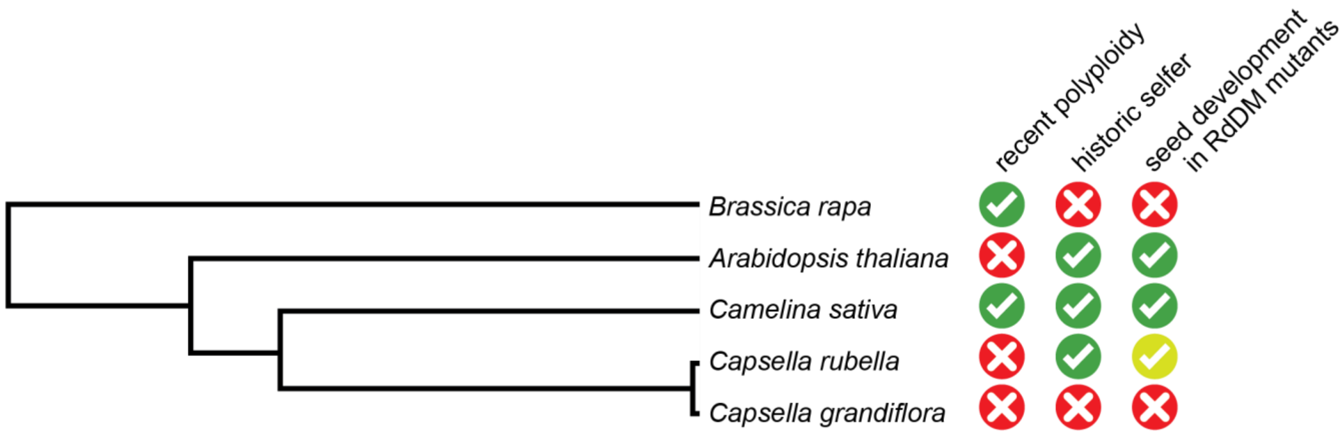


**Supplementary Figure 1. Phylogenetic relationship of study species.**
A species tree demonstrating the relationship between species in this study (based on (Nikolov et al., 2019; Brock et al., 2022)) and outlining genomic and breeding characteristics.

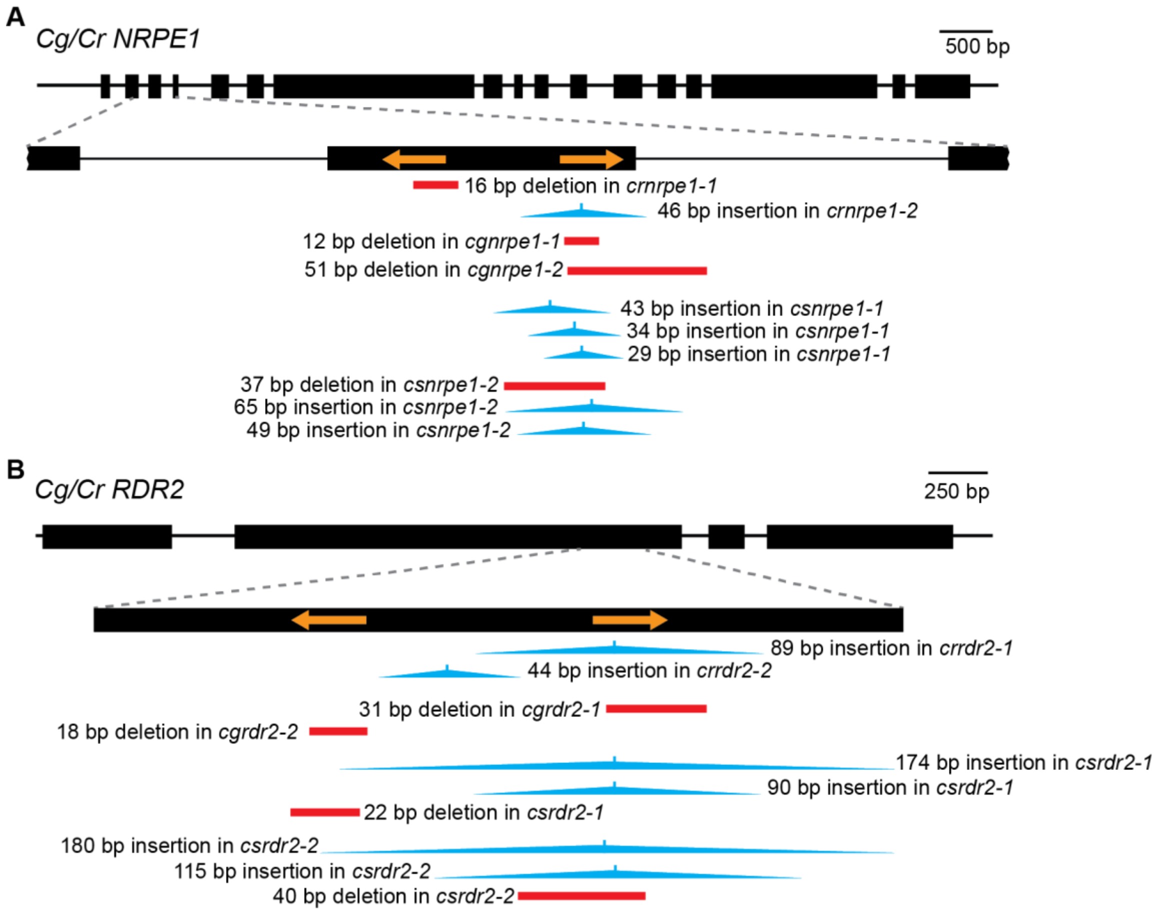


**Supplementary Figure 2. Diagram of alleles created in this study.**
*NRPE1* (A) and *RDR2* (B) gene diagrams as in Figure 1. The mutations created through genome editing are depicted below each diagram, with deletions shown by red bars and insertions depicted in blue.
